# Spatiotemporal Variation in White-Matter Development Across Early Childhood

**DOI:** 10.64898/2026.03.24.713971

**Authors:** Mervyn Singh, Dennis Dimond, Deborah Dewey, Catherine Lebel, Signe Bray

**Author notes:** Corresponding author: Mervyn Singh, PhD, Signe Bray, PhD. These authors provided equal supervisory support to this work (Catherine Lebel, Signe Bray).

## Abstract

Early childhood development is scaffolded by rapid maturation of brain white matter structure, believed to support the emergence of cognitive and socioemotional functions. Previous whole-tract studies have suggested patterns of white matter development occurring along posterior-anterior, deep-superficial and inferior-superior axes. However, little is known as to whether these patterns are evident *within* tracts. Using longitudinal diffusion imaging data from 133 children (4-8 years; 76 females), the present work characterizes along-tract patterns of white matter development across association, commissural and projection bundles using fixel-based analyses of microstructure and macrostructure. Within long range association bundles, faster age-related changes were observed for segments adjacent to the visual cortices relative to segments located near association regions, supporting a sensorimotor-association axis of brain development. An inferior-superior pattern was found for projection tracts, with faster age-effects observed for segments near the brainstem. Lastly, while several association and commissural bundles exhibited faster maturation within central segments; indicative of a deep-superficial axis, effects were mixed between micro- and macrostructure, underscoring the unique developmental timing of these different fiber properties. Our findings provide evidence that within-tract white matter maturation unfolds along key spatiotemporal axes and suggests that increased spatial precision can advance our understanding of early childhood brain development.

---

Early childhood is marked by rapid gains in cognitive, motor, and socioemotional abilities (Best and Miller 2010; Ahmed et al. 2022; Badarneh et al. 2024). These emerging skills coincide with large-scale brain maturation across distributed neural systems supporting the emergence of higher-order cognitive and behavioral functions (Nagy et al. 2004; Deoni et al. 2016; Filley and Fields 2016; Grosse Wiesmann et al. 2017; Girault et al. 2019; Bartha-Doering et al. 2021). Foundational studies using diffusion tensor imaging (DTI) have documented robust patterns in white matter micro- and macro-structural change across childhood and adolescence, and suggested that changes follow spatial developmental axes (Barnea-Goraly et al. 2005; Lebel and Deoni 2018; Tamnes et al. 2018; Lebel et al. 2019; Reynolds et al. 2019; Dimond et al. 2020; Bethlehem et al. 2022; Schilling, Chad, et al. 2023). However, given that white matter bundles traverse spatial locations, for example reaching both anterior and posterior cortical regions, ascertaining the extent of spatial patterning in white matter development requires a more spatially refined approach.

At the whole-tract level, WM development has been suggested to follow Posterior-to-Anterior (P-A), Inferior-to-Superior (I-S), and Deep-to-Superficial (D-S) axes across the brain, with posterior, ventral, and deep structures maturing earlier than anterior, superior, and superficial association pathways (Barkovich et al. 1988; Sexton et al. 2014; Simmonds et al. 2014; Lebel et al. 2019; Reynolds et al. 2019; Schilling, Archer, et al. 2023; Schilling, Chad, et al. 2023). Growing evidence further points to a Sensorimotor-Association (S-A) developmental axis as a principal organizational gradient underpinning brain development, which extends from lower-order unimodal regions (i.e., sensorimotor cortices) to transmodal association cortices (Sydnor et al. 2021; Keller et al. 2023; Sydnor et al. 2023; Luo et al. 2024). Given that individual WM pathways can span this gradient along their length, with endpoints in sensorimotor and association regions (Conner et al. 2018), research examining along-tract maturation will allow us to assess for the presence of this developmental pattern.

Along-tract techniques, where WM tracts are subdivided into discrete segments, have emerged as a powerful alternative method for mapping fine-grained spatial variations. Indeed, there is a growing body of work characterizing age-related maturation during typical development with sub-tract level resolution (Jones et al. 2005; Yeatman et al. 2012; Duerden et al. 2013; Gozdas et al. 2020; Shirazi et al. 2021; Chandio et al. 2024; Kim et al. 2025; Verschuur et al. 2025). Existing work in participants ranging from 4-30 years has demonstrated consistent posterior-anterior and inferior-superior gradients of change (Colby et al. 2011; Krogsrud et al. 2016; Lynch et al. 2020), with more recent evidence suggesting S-A and deep-superficial developmental gradients also exist (Luo et al. 2026). Longitudinal studies examining these axes across major white matter fiber bundles using advanced acquisition and processing methods are needed to build evidence for within-tract developmental patterns. Given the growing body of literature pointing to substantial along-tract variation in white matter properties, considering developmental variation within tracts is important to confirm and extend our understanding of white matter developmental axes.

Here, using high-quality longitudinal diffusion data and fixel-based analysis (FBA) (Raffelt et al. 2017; Dhollander et al. 2021), we applied an along-tract framework to map the spatiotemporal dynamics of early childhood WM development. To comprehensively map these maturational patterns across the brain, Posterior-Anterior (P-A), Deep-Superficial (D-S), Sensorimotor-Association (S-A), and Inferior-Superior (I-S) axes of development were investigated across major association (Mandonnet et al. 2018), projection (Prats-Galino et al. 2012), and commissural (Hofer and Frahm 2006) fibers. By integrating a longitudinal design with the provision of fiber-specific metrics, our study offers a precise characterization of brain structural maturation within this critical period of human development.

## Materials and Methods

### Participants

Participants were drawn from a preexisting longitudinal study investigating neurocognitive development during early childhood (Dimond et al. 2020). Children aged 4-8 years were recruited from around metropolitan Calgary and underwent a baseline MRI session and returned at 6-month, 12-month, and/or 18-month follow-up intervals. Following data processing and quality assessment, 133 children (199 scans, 76 females; age range: 4.09-7.89 years) with high-quality diffusion MRI data were included in the current study (Figure S1). Of the final sample, 73 participants (33 females) had one scan, 54 (37 females) had two scans, and 6 females had three scans. A summary of the demographic data can be found in Table S1. All participants included were born ≥ 37 weeks’ gestation, considered typically developing, with no history of neurological or psychiatric conditions. Participants were screened for contraindications to MRI scanning prior to each visit. Parents and children provided informed consent and assent before participation. Procedures were approved by the Conjoint Health and Research Ethics Board at the University of Calgary (REB14-1089).

### MRI Acquisition

All scanning sessions were performed at the Alberta Children’s Hospital. Participants underwent a mock MRI scan in an MRI simulator in the week prior to the actual assessment date to familiarize them with the scanning environment. Participants were also asked to practice lying still at home for up to 18 minutes to mitigate in-scanner movement during the MRI session. During the MRI scanning session, padding was inserted on both sides of the participant’s head to further limit any in-scanner head movement. At each visit, participants were scanned on a 32-channel 3T General Electric (GE) MR750w system (Waukesha, WI).

#### T1-weighted imaging

Whole-brain anatomical T1-weighted (T1w) images were obtained using a 3D BRAVO sequence in the sagittal plane: 0.8mm^3^ isotropic voxels, 24 x 24cm Field of View (FOV), Inversion Time (TI) = 600ms, flip angle = 10°.

#### Diffusion-weighted imaging

Diffusion MRI data was acquired in the axial plane in the anterior–posterior phase encoding direction using a 2D spin-echo Echo Planar Imaging (EPI) sequence: *b*1000 s/mm^2^ (45 directions), *b* 2000 s/mm^2^ (45 directions), and *b*0 s/mm2 (3 images); 2.5mm^3^ isotropic voxels, 45 slices, 23 x 23cm FOV, Echo Time (TE) = 86.2 msec, Repetition Time (TR) = 10 sec. Diffusion-weighted shells below *b* 1000 s/mm^2^ are susceptible to partial volume effects and can therefore bias estimates of fixel based analysis (Dhollander et al. 2021). As such, we only included the *b* 2000 s/mm^2^ shell, along with 3 *b*0 images in the present study.

### T1w Processing

T1w structural data were bias-field corrected and processed independently using the default cross-sectional reconstruction pipeline within the Computational Anatomy Toolbox, version 12 (CAT12; https://neuro-jena.github.io/cat/) (Gaser et al. 2023). One participant was missing a T1w image and was removed from the final sample. For the remaining 133 participants, estimates of Total Intracranial Volume (ICV) were then extracted for use as a covariate in statistical analysis.

### Diffusion MRI Processing

#### Pre-processing

Diffusion MRI data were processed using FSL (Jenkinson et al. 2012), ANTs (Avants et al. 2009), and MRtrix3Tissue, a fork of the MRtrix3 software package (https://3tissue.github.io/) (Tournier et al. 2019). FSL EDDY was used to perform eddy current distortion correction (Andersson and Sotiropoulos 2016) and to mitigate head motion through slice-wise outlier interpolation (Andersson et al. 2016) and motion censoring.Residual motion effects were mitigated through data censoring: volumes that had signal dropout in >20% of slices were removed, and scans were excluded from analysis if >10% of volumes were removed. The total number of slices classified as outliers for each participant (total dropout slices [TDS]) was calculated and included in our statistical analysis as a measure of head motion. The preprocessed data were then bias field corrected and spatially resampled to a voxel size of 1.25 mm isotropic to increase anatomical contrast and improve downstream model fitting and tractography.

#### Single-shell 3-Tissue CSD & Fixel-based Analysis

Tissue-specific response functions were estimated from grey matter, WM, and cerebrospinal fluid voxels in each participant’s preprocessed image, and Single-shell 3-tissue Constrained Spherical Deconvolution (SS3T-CSD) was performed to model WM fiber orientation distributions (FODs) (Dhollander and Connelly 2016). All participant FOD maps were intensity-normalized to ensure FOD amplitudes were comparable across participants and visually assessed to confirm the appropriateness of the SS3T-CSD fitting approach in resolving crossing fibers. Following model-fitting, a study-specific population template was then generated using a representative subset of 40 unique participants via linear and nonlinear registration. All participants’ FOD maps (199 scans) were subsequently warped to this population template. Fixel-based analysis (FBA), an advanced diffusion modelling approach that enables estimation of fiber bundle-specific changes in WM properties, was then implemented in template space to extract whole-brain metric maps of fiber density (FD), a microstructural measure reflecting intra-axonal volume, and fiber cross-section (FC), a morphological measure indexing macroscopic changes in fiber bundle size or shape (Raffelt et al. 2017; Dhollander et al. 2021), was computed for each participant. As per the recommended pipeline for FBA, log-transformation of FC values was then performed (FC_log_) to ensure that FC values were centered around zero and normally distributed.

#### Bundle-specific tractography

Major WM tracts of interest were delineated in population-template space using TractSeg (https://github.com/MIC-DKFZ/TractSeg), an automated tractography pipeline that provides a robust balance between the accuracy of manual dissection and the objectivity of atlas-based fiber tracking methods (Wasserthal, P. Neher, et al. 2018). As per the TractSeg workflow, bundle-specific binary voxel masks and track-oriented maps (TOM) (Wasserthal, P.F. Neher, et al. 2018), were created for each tract using the peak FOD values derived from the population template. These precursor files were then used as inputs for probabilistic tractography to generate 10,000 streamlines per tract in each hemisphere via the default second-order integration over fiber orientation distribution (iFOD2) algorithm (Wasserthal et al. 2019).

We chose to delineate 17 WM tracts of interest spanning the three major bundle classes: (i) ***Association tracts:*** Arcuate fasciculus [AF], Cingulum [CG]; Inferior-fronto-occipital fasciculus [IFO]; Inferior-longitudinal fasciculus [ILF]; Segments 1-3 of the Superior-longitudinal fasciculus [SLFI-III], (ii) ***Projection tracts*:** Corticospinal tract [CST]; Fronto-pontine tract [FPT]; Parieto-occipital-pontine tract [POPT]), and lastly, (iii) ***Commissural tracts:*** Subdivisions 1-7 of the Corpus callosum: Rostrum (CC-1), Genu (CC-2), Premotor (CC-3), Primary motor (CC-4), Primary somatosensory (CC-5), Primary isthmus (CC-6) and Splenium (CC-7). All tracts were then visualized on the population template to ensure each was appropriately delineated.

#### Along-tract Segmentation

Along-tract regions of interest were defined for each tract using a combination of data-driven and standardized approaches. First, QuickBundles (Garyfallidis et al. 2012), a Python-based tract clustering algorithm, was used to identify centroids, representing the central streamline of each tract. Each tract was then subdivided into 20 equidistant segments along its centroid using a custom script in MATLAB (Version R2024b) that resampled individual streamline points to their nearest centroid segment. Segments were ordered systematically: association tracts from anterior (Segment 1) to posterior (Segment 20), projection tracts from superior (Segment 1) to inferior (Segment 20), and commissural tracts from the left (Segment 1) to right (Segment 20) hemisphere. Streamline points belonging to each segment were subsequently mapped onto the whole brain fixel maps for each participant and cropped to generate regions of interest containing only those fixels lying within the anatomical boundaries of each segment. A visual representation of the along-tract segmentation workflow for three representative tracts (***Association*:** IFO, ***Projection*:** CST, ***Commissural*:** CC-4) is provided in Figure 1.

**Fig. 1.**
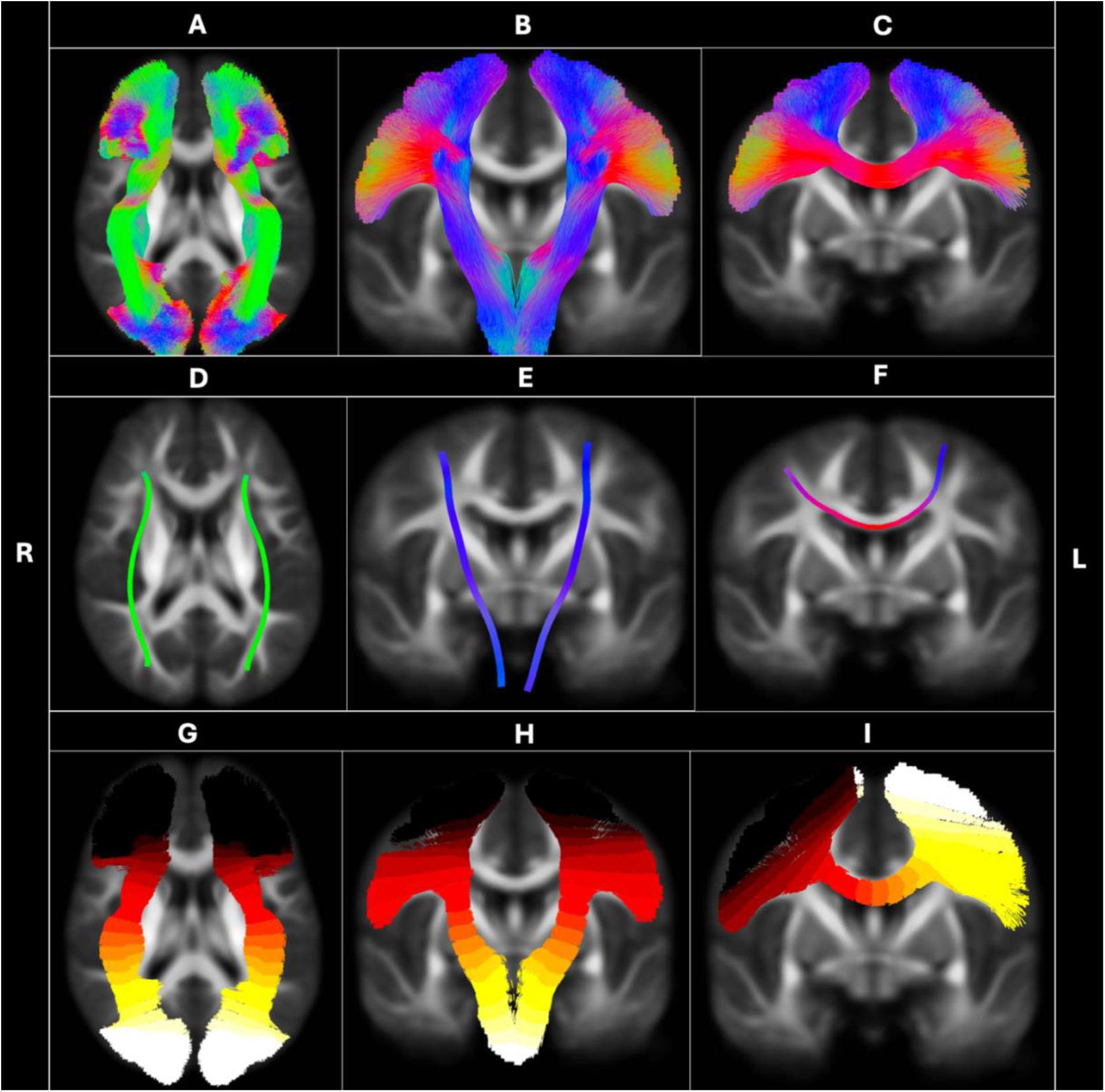
Representation of the along tract segmentation workflow for a subset of tracts. Top panel (A, B, C): streamlines for the bilateral IFO, CST and CC-4. Middle panel (D, E, F): centroids for the bilateral IFO, CST and CC-4. Bottom panel (G, H, I): ROI segments for the bilateral IFO, CST and CC-4 Tract streamlines and centroids are coloured according to their principal orientation (inferior-superior: blue; anterior-posterior: green; transversal: red). Tract segments are coloured using the ‘hot’ colour palette. Segments are colored in ascending order from dark to light.

#### Quality Assessment

Segment-wise fixel masks were threshholded to minimize the influence of spurious FD and FC_log_ values. This was undertaken across several stages. First, for each tract segment, fixels with FD values in the bottom 25th percentile were identified for all subjects individually, and the highest value across all participants was selected as the cutoff (see Supplementary Figures S2-S4 for line plots visualizing the segment-level FD thresholding criteria for each segment). Fixels below this threshold were then removed from segment masks across all participants. We then applied the threshholded FD maps to their corresponding FC_log_ maps to remove all non-overlapping FC log fixels, thus ensuring spatial consistency of all fixels across both metric maps for that tract segment. Following segment-level thresholding, Segments 18-20 were removed from the bilateral CST, FPT, and POPT tracts due to field-of-view cropping artefacts in the inferior regions of the brain across many participants, while segments 1-3 were excluded from the bilateral ILF due to low fixel retention after thresholding, and cropping artefacts in anterior portions. In addition, we excluded any remaining cases with less than 75% retained fixels for a given tract segment (3,635 individual tract segments), resulting in a final sample of 99,565 tract segments. Lastly, average values of FD and FC_log_ were calculated within each segment-specific fixel mask to include as outcome measures in our statistical models.

### Statistical Analyses

Statistical analyses were performed in R (version 4.4.2, Core Team, 2023). To examine spatiotemporal patterns of WM maturation in early childhood, we conducted our analyses across the following two levels of investigation:

### Segment-wise developmental changes in WM

Along-tract profiles of WM development were assessed by conducting a series of linear-mixed effects (LME) models (Brown 2021; da Silveira et al. 2023) using the lme4 package (version 1.1-36) (Bates et al. 2014) predicting mean FD or FC_log_ from age within each tract segment. All models included sex assigned at birth (female, male), handedness (right, left, or ambidextrous), intracranial volume (ICV), and total number of slices identified as being outliers (i.e., “total dropout slices”; TDS) as covariates of no interest. Continuous predictors were standardized prior to running all models to allow for comparisons to be made across variables. To account for participant-level clustering, participant ID was included as a random effect in all models. Correction for multiple comparisons was performed across all segments within each tract using the False Discovery Rate (FDR) method (Benjamini and Hochberg 1995), resulting in 20 comparisons per tract in each hemisphere. Significance was set to an adjusted alpha value of p_adj_ <.05. Standardized Rates of change for FD and FC_log_ (beta-coefficients; β^) were extracted and plotted for each tract segment to visualize age-related effects in WM micro- and macrostructure.

### Spatiotemporal axes of WM maturation

Spatiotemporal axes of age-related change along each tract were assessed using linear regression analyses. Standardized beta-coefficients (β^) derived from the aforementioned LME models that estimated the association between age and segment-wise measures of FD and FC_log_ were extracted to determine whether these age-related effects varied systematically according to segment location along the following principal developmental axes: (i) **posterior-anterior (P-A**)*, (ii)* **sensorimotor-association (S-A),** *(iii)* **inferior-superior (I-S)** and (iv) ***deep-superficial (D–S)*.**

#### Association tracts

For association tracts (AF, CG, SLF-I – III), both P-A and D-S axes were modelled. To assess change in age effects along the P-A axis, segment locations were coded as continuous values from 1 (anterior) to 20 (posterior). For the D–S axis, segment locations were re-labelled symmetrically from –9.5 to 9.5 and converted to absolute values, such that increasing values represented greater distance from the tract midpoint, allowing assessment of variation in age effects with distance from the midline. For a subset of long-range association tracts with posterior segments reaching the occipital cortex (IFO, ILF), the P-A direction also reflects variation along the S-A axis (Sydnor et al. 2021). To assess age changes along the S-A axis, segment location was coded continuously in ascending order from sensorimotor (1) to association (20).

#### Projection tracts

For projection tracts (CST, FPT, POPT), segment locations were relabeled in ascending order from ‘inferior’ (subcortical and brainstem adjacent segments) to ‘superior’ (segments extending towards the cortex) to examine maturation along the I-S axis.

#### Commissural tracts

For commissural tracts (CC-1 through CC-7), only the D–S term was modeled, with segment location recoded symmetrically following the same approach as implemented for the association tracts.

#### Statistical Interpretation

For P-A and S-A axes, negative beta-coefficients indicate faster maturation towards the anterior or association cortices, whilst positive coefficients represent faster maturation towards the posterior or sensorimotor cortices. For the I-S axis, negative coefficients represent faster maturation towards inferior segments, and positive coefficients indicate faster maturation towards the superior. Lastly, for the D-S axis, negative values represent faster maturation towards superficial segments of a tract, whilst positive values suggest faster maturation towards deeper segments. Correction for multiple comparisons was conducted using the Bonferroni method within each Tract-Group (Association, Projection, Commissural) * Axis (P-A/S-A, D-S, I-S) * Metric (FD, FC_log_) combination.

## Results

### Association tracts

#### Posterior-Anterior Axis

Spatiotemporal patterns of FD and FC_log_ along the Posterior-Anterior (P-A) axis are presented in Figure 2, Panel C. Evidence of faster FD maturation was found in posterior segments within the bilateral CG, SLF-I and right SLF-III. In contrast, faster maturation toward the anterior end was found for the bilateral SLF-II (left: β^= -0.155; right: β^ = -0.24) and left SLF-III (β^ = -0.167). For FC_log_, most association tracts demonstrated faster maturation within anterior segments, with the strongest effects evident in the bilateral SLF-I (left: β^ = -0.922; right: β^ = -0.466) and SLF-III (left: β^ = -0.243; right: β^ = -0.332).

**Fig. 2.**
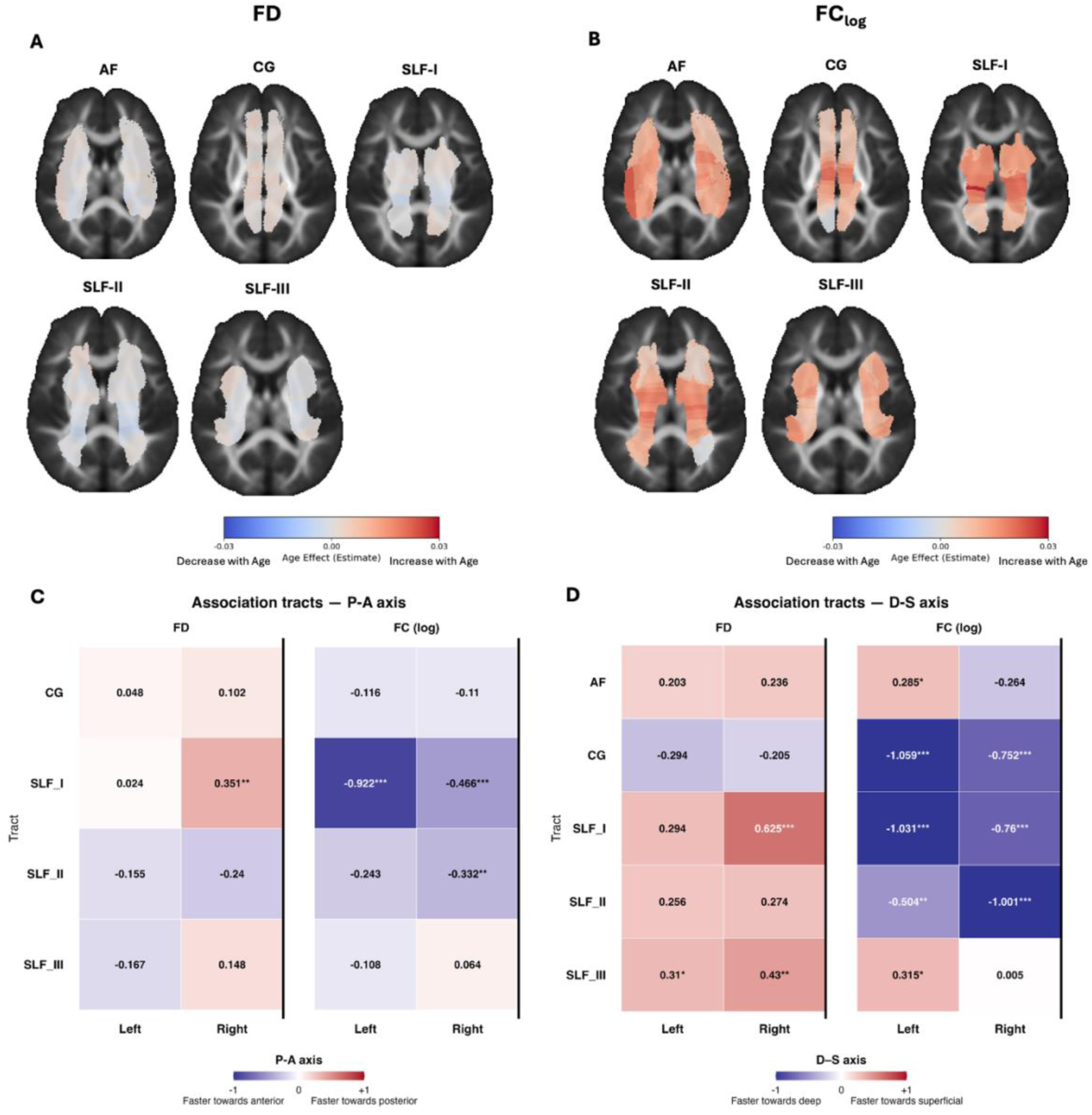
Posterior-Anterior (P-A) and Deep-Superficial (D-S) age-related changes in along-tract WM properties within association tracts. *Top Panels:* Segment-wise age-related changes in fiber density (FD; Panel A, left) and log-transformed fiber cross-section (FC_log_; Panel B, right) in the bilateral Arcuate fasciculus (AF), Cingulate gyrus (CG), and the Superior longitudinal fasciculus I-III (SLF-I, SLF-II, SLF-III). Tract segments are color-coded by their corresponding standardized age effect size estimates derived from linear mixed models (beta-coefficients) and overlaid onto an axial slice of the population template. Warmer colors indicate greater age-related increases in WM properties. Cooler colors indicate greater age-related decline. *Bottom Panels:* Heatmap plots illustrating spatiotemporal patterns of FD and FC_log_ along the P-A (Panel C) and D–S (Panel D) axes. Values represent the strength and directional bias of maturation effects along the axis. Positive values and warmer colors reflect stronger age-effects in posterior (P-A) and superficial (D–S) segments of the tracts, whereas cooler colors reflect stronger age-effects in anterior (P-A) and deeper (D–S) segments of the tract. Statistical significance is denoted as: \**p* < 0.05, \*\**p* < 0.001, \*\*\**p* < 0.0001.

#### Deep-Superficial Axis

Spatiotemporal patterns of FD and FC_log_ along the Deep-Superficial (D-S) axis are presented in Figures 2 and 3, Panel D. Most association tracts showed faster maturation in FD within superficial segments, except for the bilateral CG (left: β^ = -0.294; right: β^ = -0.205), which demonstrated faster maturation in deeper segments. For FC_log_, most tracts exhibited faster maturation in deeper segments, particularly in the right hemisphere. In contrast, the bilateral ILF showed strong age-related maturation within superficial segments (left: β^ = 0.899; right: β^ = 0.677).

**Fig. 3.**
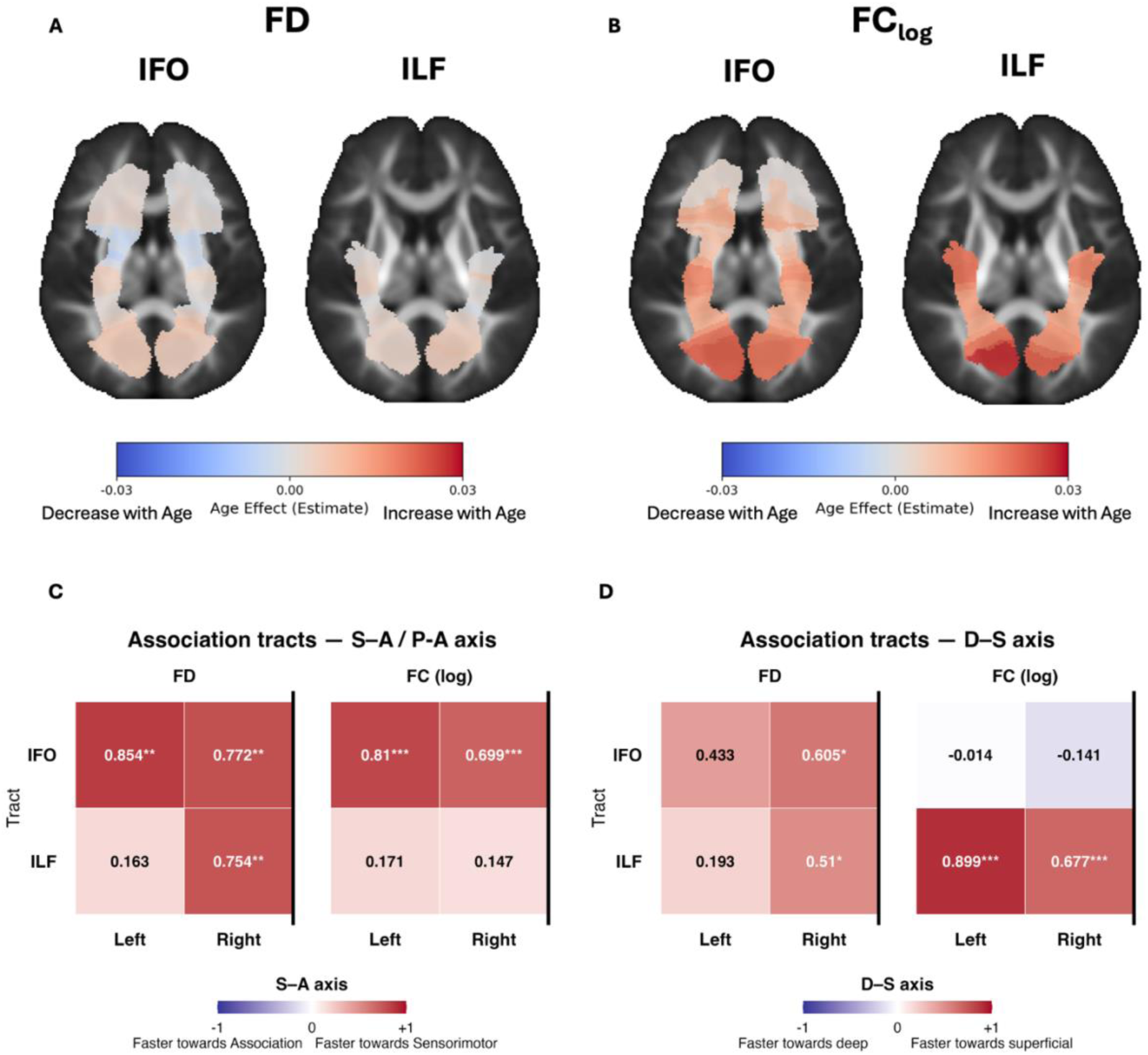
Sensorimotor-Association (S-A)/Posterior-Anterior (P-A) and Deep-Superficial (D-S) age-related changes in along-tract WM properties within long range association tracts. *Top Panels:* Segment-wise age-related changes in fiber density (FD; Panel A, left) and log-transformed fiber cross-section (FC_log_; Panel C, right) in the bilateral Inferior fronto-occipital fasciculus (IFO) and Inferior longitudinal fasciculus (ILF), Tract segments are color-coded by their corresponding standardized age effect size estimates derived from linear mixed models (beta-coefficients) and overlaid onto an axial slice of the population template. Warmer colors indicate greater age-related increases in WM properties. Cooler colors indicate greater age-related decline. *Bottom Panels:* Heatmap plots illustrating spatiotemporal patterns of FD and FC_log_ along the S-A (Panel C) and D-S (Panel D) axes. Values represent the strength and directional bias of age effects along the axes. Positive values and warmer colors reflect stronger age-effects in sensorimotor (S-A) or superficial (D-S) adjacent segments, whereas cooler colors reflect stronger age-effects in association (S-A) or deeper (D-S) segments. Statistical significance is denoted as: \**p* < 0.05, \*\**p* < 0.001, \*\*\**p* < 0.0001.

#### Sensorimotor-Association Axis

Spatiotemporal patterns of FD and FC_log_ along the Sensorimotor-Association (S-A) axis for long range association fibers (IFO and ILF) are presented in Figure 3, Panel C. Results demonstrate faster maturation in both FD and FC_log_ in segments proximal to the sensorimotor cortices compared to segments closer to the association regions.

### Projection tracts

Segment-wise age-related changes in FD and FC_log_ in projection tracts (CST, POPT, FPT) are presented in Figure 4 (Panel A). Spatiotemporal patterns of FD and FC_log_ along the Inferior–Superior (I-S) axis are presented in Panel B. All projection tracts showed consistent spatiotemporal patterns along the I-S axis, with faster maturation in FD and FC_log_ in the inferior segments. The strongest effects were observed for FD in the right CST (β^ = –0.642) and right FPT (β^ = –0.757), and for FC_log_ in the right FPT (β^ = –0.712) and bilateral POPT (left: β^ = –0.828; right: β^ = –0.673).

**Fig. 4.**
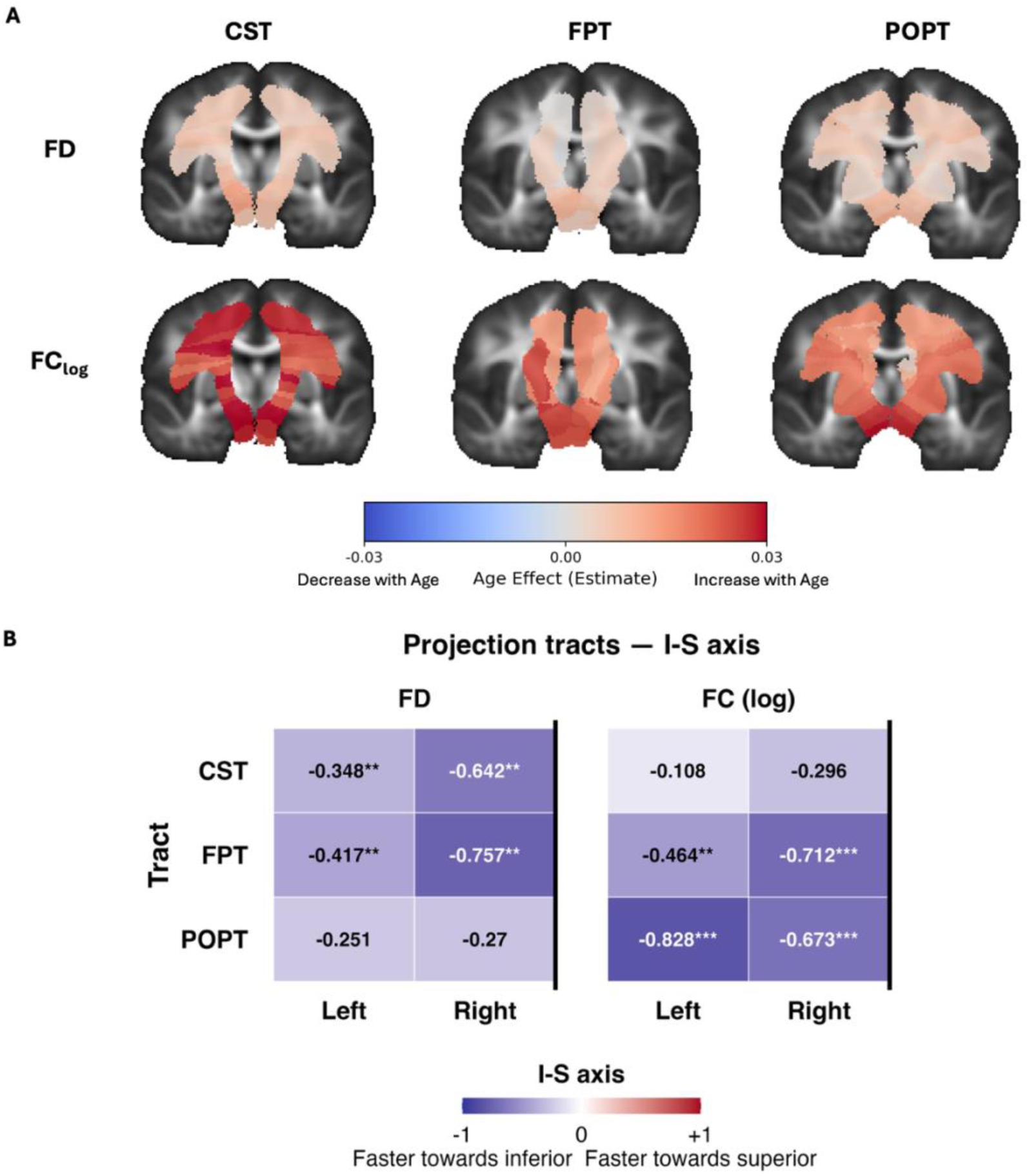
Inferior-Superior (I-S) age-related changes in along-tract WM properties within projection tracts. *Top Panels:* Segment-wise age-related changes in fiber density (FD; Panel A top) and log-transformed fiber cross-section (FC_log_; Panel A bottom) in the bilateral Corticospinal tract (CST), Fronto-pontine tract (FPT) and Parieto-occipital-pontine (POPT). Tract segments are color-coded by their corresponding standardized age effect size estimates derived from linear mixed models (beta-coefficients) and overlaid onto a coronal slice of the population template. Warmer colors indicate greater age-related increases in WM properties. Cooler colors indicate greater age-related decline. *Bottom Panels:* Heatmap plots illustrating spatiotemporal patterns of FD and FC_log_ along the I-S axis (Panel B). Values represent the strength and directional bias of age effects along the axis. Positive values and warmer colors reflect stronger age-effects in superior segments, whereas negative values and cooler colors reflect stronger age-effects in inferior segments. Statistical significance is denoted as: \**p* < 0.05, \*\**p* < 0.001, \*\*\**p* < 0.0001.

### Commissural tracts

Segment-wise age-related changes in FD and FC_log_ in the seven major subdivisions of the corpus callosum (CC-1 – CC-7) are presented in Figure 5 (Panel A & B). Spatiotemporal patterns of FD and FC_log_ along the Deep–Superficial (D-S) axis are presented in Panel D. Faster FD maturation in superficial segments was evident across all corpus callosum subdivisions, with the strongest effects observed in CC-1 (β^ = 0.392) and CC-7 (β^ = 0.358). For FC_log_, while most callosal subdivisions showed faster maturation within superficial segments, the CC-2 (β^ = –0.640) and CC-6 (β^ = –0.233), showed faster maturation toward deeper segments.

**Fig 5.**
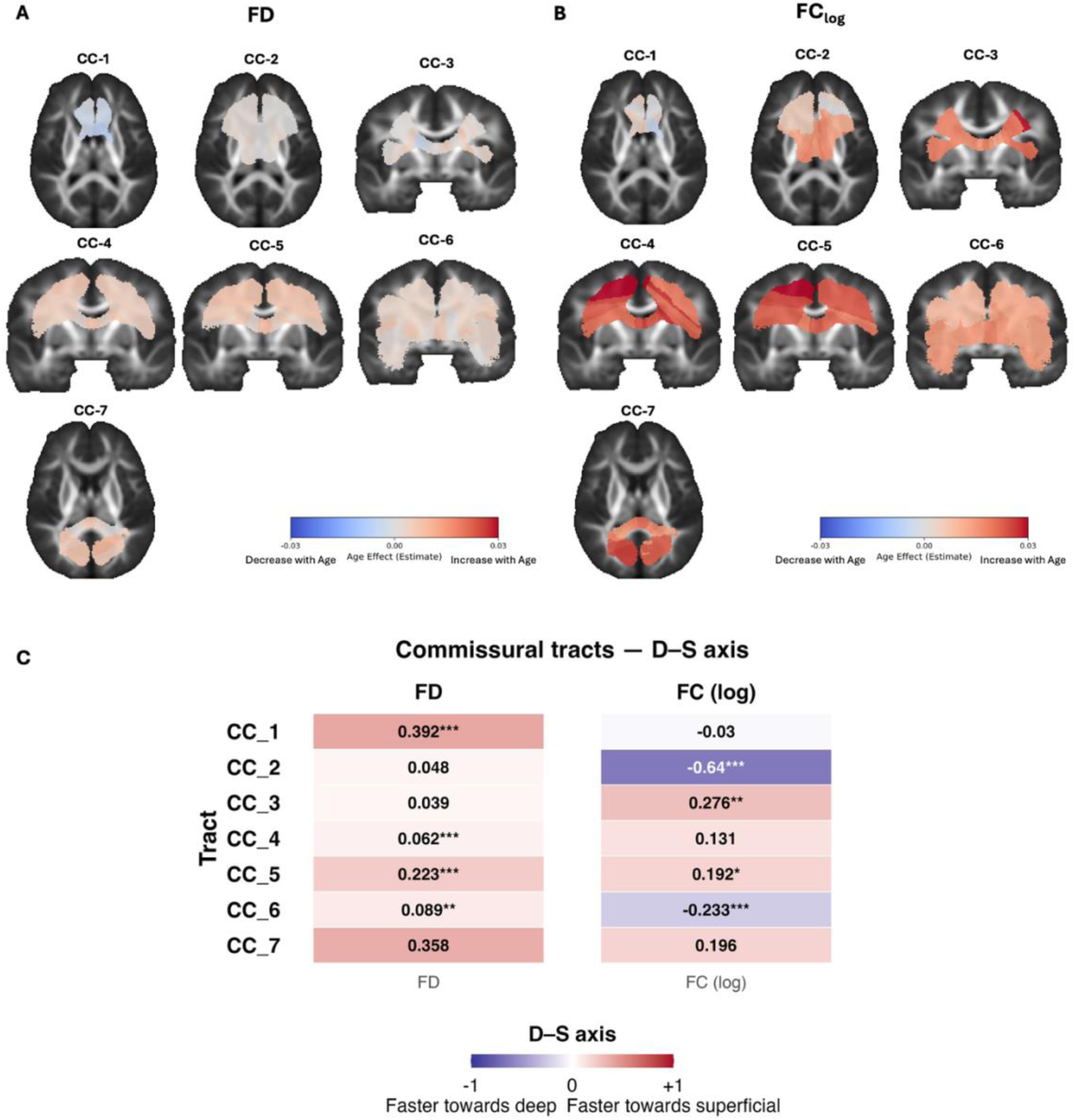
Deep-Superficial (D-S) age-related changes in along-tract WM properties within the seven major subdivisions of the Corpus Callosum (CC). *Top Panels:* Segment-wise age-related changes in fiber density (FD; Panel A, left) and log-transformed fiber cross-section (FC_log_; Panel B, right) in the Rostrum (CC-1), Genu (CC-2), Premotor (CC-3), Primary motor (CC-4), Primary somatosensory (CC-5), Primary isthmus (CC-6) and Splenium (CC-7). Tract segments are color-coded by their corresponding standardized age effect size estimates derived from linear mixed models (beta-coefficients) and overlaid onto a coronal slice of the population template. Warmer colors indicate greater age-related increases in WM properties. Cooler colors indicate greater age-related decline. *Bottom Panels:* Heatmap plots illustrating spatiotemporal patterns of FD and FC_log_ along the D–S (Panel D) axes. Values represent the strength and directional bias of age effects along the axis. Positive values and warmer colors reflect stronger age-effects in superficial segments, whereas negative values and cooler colors reflect stronger age-effects in deeper segments. Statistical significance is denoted as: \**p* < 0.05, \*\**p* < 0.001, \*\*\**p* < 0.0001.

## Discussion

The present work implemented an along-tract, fixel-based analysis to characterize age-related changes in along-tract WM development across 17 major association, commissural and projection tracts in children aged 4-8 years. We observed several distinct spatiotemporal patterns across the three major bundle classes. First, in association bundles, tracts with projections in the visual cortex exhibited a clear spatiotemporal pattern along the S-A axis, while P-A effects were more heterogenous across tracts that did not terminate in the visual areas. Second, while there was some evidence of a deep-to-superficial axis across association and commissural bundles, the direction of these effects was mixed and differed between micro- and macro-structure. Lastly, we found support for an inferior-to-superior pattern for projection tracts connecting the motor cortices to subcortical structures. These findings underscore the importance of considering within-tract variation when examining WM maturation and provide new insights into WM changes during an important period of childhood development.

Within long-range association bundles (IFO, ILF), we observed stronger age-related rates of change of both microstructural (FD) and macrostructural (FC_log_) properties in segments proximal to sensorimotor cortices relative to those near higher-order association regions. These findings converge with those of Luo et al. (2026) and provide further support that WM development follows a sensorimotor-association (S-A) axis not only across, but within tracts. The cortical hierarchy along the S-A axis describes brain development progressing from evolutionarily older sensorimotor areas, which are primarily unimodal in functionality, toward transmodal association regions that demonstrate prolonged plasticity and extended developmental timelines due to their role in higher-order cognition (Yakovlev 1967; Deoni et al. 2015; Grydeland et al. 2019; Dong et al. 2021; de Faria et al. 2021; Sydnor et al. 2021; Valk et al. 2022; Gao 2025). The IFO and ILF link cortical regions at divergent ends of the S-A continuum, and our findings align with this organizational gradient, as segments adjacent to the visual cortices appear to undergo accelerated change in early childhood, whereas weaker age-effects in segments situated near the association cortices may be indicative of the protracted maturation patterns in these regions. In contrast to the clear S-A patterns evident in long range association tracts, spatiotemporal effects were more varied in P-A oriented pathways that did not terminate in the visual areas (i.e. SLF-I - III; CG). While FD demonstrated faster maturation in posterior segments, FC_log_ showed the opposite pattern of faster age-effects towards anterior segments. This dissociation in spatiotemporal patterns suggests that micro- and macrostructural properties follow partially independent spatiotemporal patterns within the same tract in early childhood.

Our investigation of a deep-to-superficial (D-S) axis of maturation across association and commissural fibers revealed mixed findings. Faster age-related effects for FC_log_ were observed within deep WM segments across most association tracts, whereas FD showed the opposite pattern, with relatively faster maturation in superficial segments closer to the cortical mantle. For commissural tracts, most callosal regions exhibited faster FD and FC _log_ maturation in segments nearer to the cortical surface, mirroring the patterns observed in association tracts. Faster FC_log_ effects in deeper association tract segments align with evidence from small-scale studies of infants, demonstrating (Gao et al. 2009; Williamson and Lyons 2018; Grotheer et al. 2022). Indeed, superficial WM fibers are often the last to mature in adulthood (Guevara et al. 2020; Schilling, Archer, et al. 2023; Zhang et al. 2024). It is important to note that Luo et al. (2026) reported less pronounced age-related changes in core segments compared to superficial regions, reflecting stabilization of maturational effects later in development. Given the relatively young age range of children in our cohort (4-8 years), our observation of faster age-related FC_log_ effects likely reflects ongoing maturation in these core regions during early childhood, which have not yet stabilized. We speculate that as children transition into later childhood and adolescence, age-related effects in deep segments may attenuate (as seen in Luo et al., 2026), giving way to accelerated maturation within superficial segments.

Lastly, projection tracts exhibited highly consistent patterns that unfold along an inferior-to-superior (I-S) axis. That is, greater age-related increases in FD and FC_log_ were observed in inferior segments, compared with weaker rates of change in segments closer to the sensorimotor cortices. Inferior-to-superior patterns have previously been identified in post-mortem myelin staining work (Yakovlev 1967), and in the limited along-tract literature (Colby et al. 2011; Sexton et al. 2014; Krogsrud et al. 2016). Our results suggest that inferior segments of projection pathways, which connect subcortical nuclei, undergo congruent microstructural and macrostructural refinement during early childhood, supporting the emergence of motor coordination and sensory feedback functions (Kamali et al. 2010; Prats-Galino et al. 2012; Welniarz et al. 2017; Florio 2025). Similar to the D-S maturational profiles seen above, we speculate that age-effects within inferior segments would reach a maturational peak and attenuate during later childhood and adolescence, giving way to more prolonged changes in cortical WM properties.

Our results add to a growing literature demonstrating that maturational processes are highly heterogeneous within individual fiber bundles and are the first to characterize spatially varying patterns of WM development within tracts using a fixel-based analysis framework. By employing fixel-based metrics, we were able to uncover unique developmental patterns for micro- and macrostructure. Across all investigated tracts, we found that segment-level macrostructural properties exhibited consistently stronger age-related effects than microstructural properties. This overall pattern is largely in agreement with prior whole-tract fixel-based analysis studies (Genc et al. 2018; Dimond et al. 2020), suggesting that macrostructural remodeling may represent a key mechanism underlying WM maturation during the early childhood developmental window. The more limited age-related effects observed for FD may reflect earlier stabilization of microstructural organization. Alternatively, age effects may emerge during later childhood and adolescence following a relative attenuation of large-scale morphological change. Future longitudinal studies spanning a broader developmental range will be necessary to test directly whether the balance between macrostructural expansion and microstructural refinement shifts spatially across distinct phases of WM maturation. Whilst the present study is strengthened by focusing on an important and understudied developmental period and the inclusion of longitudinal data, we note several limitations. First, the sample size may be underpowered for detecting small segment-level changes with multiple comparisons correction, limiting interpretation of individual segment-level effects. Second, despite the increased specificity offered by fixel-derived measures, these metrics remain indirect proxies for WM organization. For example, higher FD values, reflecting increased intra-axonal compartment diffusion signal may reflect increased axonal packing, calibre, and/or myelination (Raffelt et al. 2012; Dhollander et al. 2021). Future studies should adopt a multimodal approach, inclusive of myelin-sensitive imaging techniques (Varma et al. 2015; Geeraert et al. 2019; Munsch et al. 2021), to better disentangle the underlying biological mechanisms driving spatial variability in within-tract maturation.

## Conclusions

In summary, we have demonstrated that WM development during early childhood unfolds heterogeneously within tracts. Through the implementation of fixel-based analysis, our findings reinforce the presence of sensorimotor-association, deep-superficial and inferior-superior axes within specific bundle groups. Collectively, our work underscores the importance of considering within-tract variability in white matter maturation and provides a more nuanced framework for understanding typical brain development during a critical period of early childhood development.

## Author Contributions

**Mervyn Singh –** Conceptualization, Methodology, Formal analysis, Visualization, Writing - Original Draft, Writing - Review & Editing. **Dennis Dimond –** Conceptualization, Methodology, Formal analysis, Writing - Review & Editing. **Deborah Dewey –** Writing - Review & Editing. **Catherine Lebel –** Conceptualization, Methodology, Writing - Original Draft, Writing - Review & Editing, Supervision. **Signe Bray –** Conceptualization, Methodology, Resources, Investigation, Writing - Original Draft, Writing - Review & Editing, Supervision, Project administration, Funding acquisition.

## Acknowledgments

Computational resources and infrastructure for processing and analysis of neuroimaging data were provided by the University of Calgary Advanced Research Computing (ARC) cluster. We thank the research staff at the Alberta Children’s Hospital Research Institute, Hotchkiss Brain Institute, and the Owerko Centre for Neurodevelopment & Child Mental Health for their assistance with the study. Finally, we are grateful to all the parents, guardians, and children who participated in this study.

## Data availability statement

Due to ethical considerations, public sharing of the raw data is not permitted but may be provided upon reasonable request by contacting the senior corresponding author (S. Bray). This policy is in accordance with the funding bodies that supported this research and the Conjoint Health and Research Ethics Board at the University of Calgary. Scripts for all steps in the analysis pipeline are publicly available on GitHub: https://github.com/MervSingh/alongtract_fba_wm_dev.git

