## Supplementary Materials for "Spatiotemporal Variation in White-Matter Development Across Early Childhood"

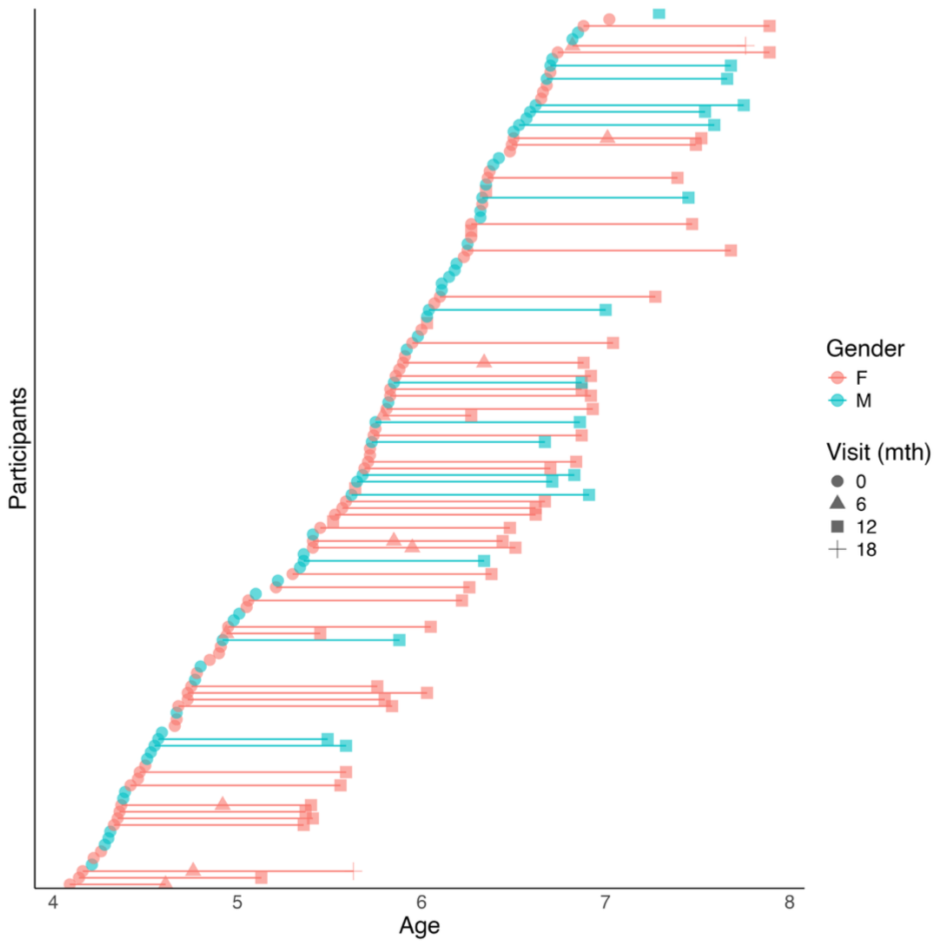


**Fig S1.** Spread of participants by age. Each row depicts a single participant, and each data point represents a single scan (females = pink; males = blue). Lines connecting each data point indicate multiple scans by the same participant.

**Table S1.** Sample characteristics at each timepoint.

| | Baseline | 6-month | 12-month | 18-month |
|---|---|---|---|---|
| n | 125 | 10 | 63 | 2 |
| Age (years), M (SD) | 5.55 (0.82) | 5.7 (0.86) | 6.57 (0.76) | 6.7 (1.51) |
| [range] | [4.09-7.02] | [4.61-7.01] | [5.13-7.89] | [5.63-7.76] |
| Sex, F/M | 68/56 | 10/0 | 45/18 | 2/0 |
| Hand, R/L/A | 69/8/0 | 10/0/0 | 57/6/0 | 2/0/0 |
| ICV, M (SD) ($mm^3$) | 973417.56 | 278476.95 | 1094225.66 | 1118643.49 |
| | (550435.45) | (583876.09) | (521656.57) | (347439.89) |
| TDS, M (SD) | 26.82 (18.86) | 10.8 (6.83) | 20.65 (13.73) | 20.5 (20.51) |

Note. F: Female; M: Male; R: Right/Mostly Right; L: Left/Mostly Left; A: Ambidextrous; ICV: Intracranial Volume; TDS: Total Number of Dropout Slices.

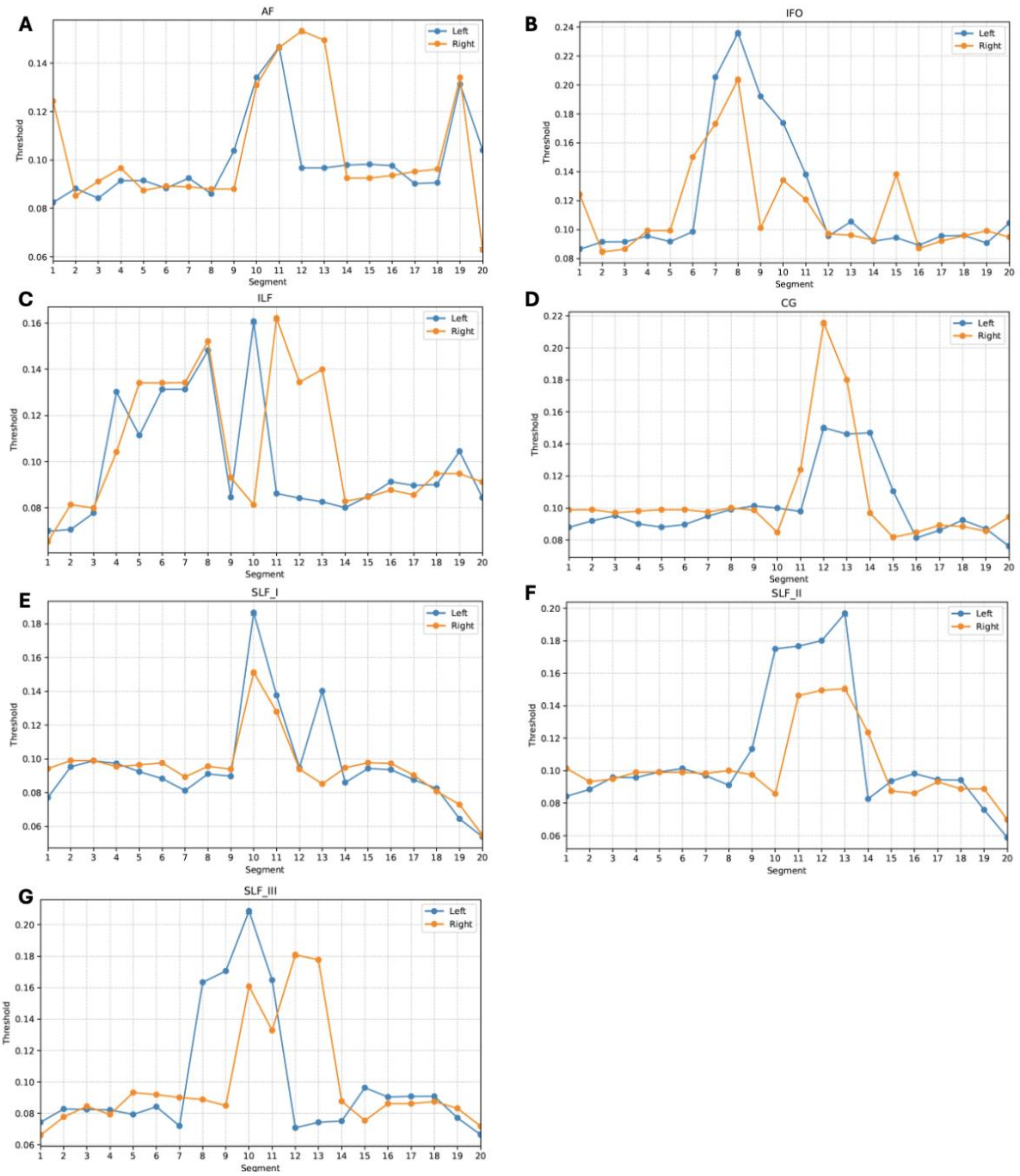


**Fig S2**. Segment-wise fixel thresholding of Association tracts. *Panels A-G:* Line plots visualizing the fiber density (FD) threshold criteria across segments of the Arcuate fasciculus (AF; Panel A), Inferior fronto-occipital fasciculus (IFO; Panel B), Inferior longitudinal fasciculus (ILF; Panel C), Cingulate gyrus (CG; Panel D), and the Superior longitudinal fasciculus I-III (SLF-I, SLF-II, SLF-III; Panels D-G). Fixels falling below these thresholds were removed from each participant's FD map. The threshholded FD maps were then applied to their corresponding $FC_{log}$ maps to ensure spatial correspondence. Blue: Threshold values across segments for the left hemisphere. Yellow: Threshold values across segments for the right hemisphere.

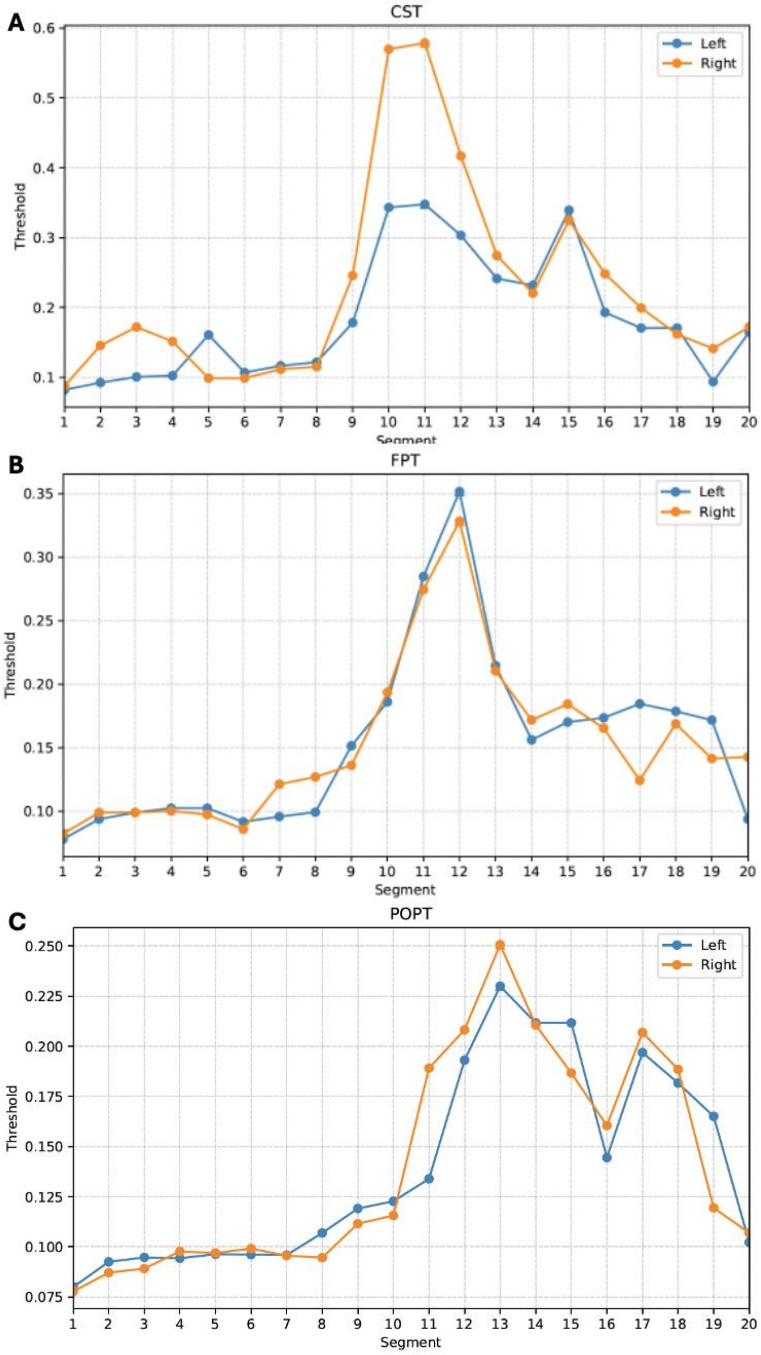

**Fig S3.** Segment-wise fixel thresholding of Projection tracts. *Panels A-C*: Line plots visualizing the fiber density (FD) threshold criteria across segments of the Corticospinal tract (CST), Fronto-pontine tract (FPT) and Parieto occipital-pontine tract (POPT). Fixels falling below these thresholds were removed from each participant's FD map. The threshholded FD maps were then applied to their corresponding $FC_{log}$ maps to ensure spatial correspondence. Blue: Threshold values across segments for the left hemisphere. Yellow: Threshold values across segments for the right hemisphere.

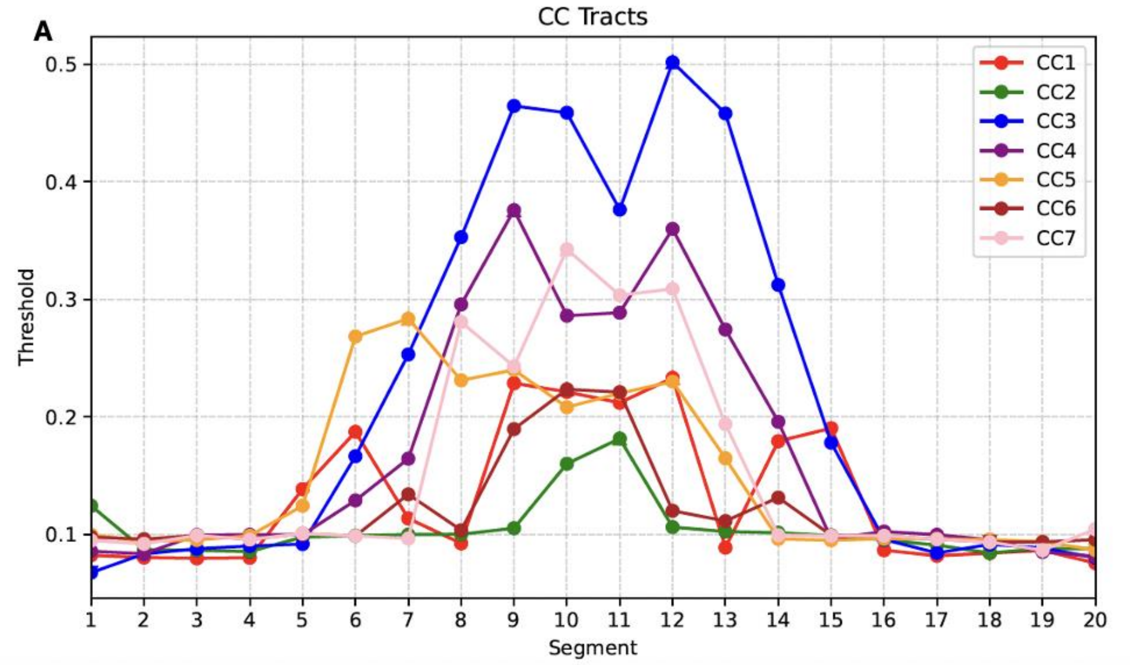


**Fig S4.** Segment-wise fixel thresholding within the seven major subdivisions of the Corpus Callosum (CC). *Panel A:* Line plots visualizing the fiber density (FD) threshold criteria across segments of the Rostrum (CC-1; Red), Genu (CC-2; Green), Premotor (CC-3; Blue), Primary motor (CC-4; Purple), Primary somatosensory (CC-5; Yellow), Primary isthmus (CC-6; Maroon) and Splenium (CC-7; Pink). Fixels falling below these thresholds were removed from each participant's FD map. The threshholded FD maps were then applied to their corresponding $FC_{log}$ maps to ensure spatial correspondence.
